# Shifts in microbial transcription patterns prior to onset of *Necrotizing enterocolitis* may indicate low oxygen levels in the gut

**DOI:** 10.1101/466417

**Authors:** Yonatan Sher, Matthew R. Olm, Tali Raveh-Sadka, Christopher Brown, Ruth Sher, Brian Firek, Robyn Baker, Michael Morowitz, Jillian F. Banfield

## Abstract

Premature infants are at risk for developing *necrotizing enterocolitis* (NEC), an inflammatory disease that can progress to necrosis of gut tissue. Previous attempts have failed to identify any consistent predictor of NEC. We hypothesized that prior to the appearance of NEC symptoms, the gut microbiome shifts in its transcriptional profile. To test this hypothesis we integrated genome-resolved metagenomic and metatranscriptomic data from multiple time-points in the first month of life of four preterm infants, two of whom later developed NEC. Gut microbiomes of NEC infants showed increased transcription of high oxygen affinity cytochrome oxidases and lower transcription of genes to detoxify nitric oxide, an antimicrobial compound released by host cells. These results, and high transcription of H_2_ production genes, suggest low O_2_ conditions prior to NEC onset, and are consistent with hypoxic conditions in diseased gut tissue. The findings motivate further testing of transcript data as a predictor of NEC.

**Highlights:**

- Transcription of high oxygen affinity microbial cytochrome oxidase may predict *necrotizing enterocolitis* (NEC) development.
- Lower transcription of microbial genes to detoxify nitric oxide (NO) may also predict NEC development.
- Higher transcription of H_2_ production genes by *Escherichia sp.* was found in the gut of premature infants that develop NEC.

---

*Necrotizing enterocolitis* (NEC) is a potentially lethal gut disease that almost exclusively strikes premature infants. The hallmark of NEC is inflammation of the small and/or large bowel that can progress rapidly to intestinal necrosis, sepsis, and death (Hackam and Caplan, 2018; Neu and Walker, 2011). Because the onset of the disease is often fulminant, treatment options for severe cases are limited and often futile. Consequently, in order to develop new ways to diagnose and treat NEC as early as possible, much of the recent research on NEC focuses on its early development stages (Niño et al., 2016).

The need for disease biomarkers to enable early and accurate diagnosis motivates the ongoing search for the cause of NEC. Aberrant gut bacterial colonization (Kliegman, 1985), or microbial dysbiosis (Warner et al., 2016), has been considered to be an important risk factor for NEC. Evidence supporting the role of bacteria in NEC includes positive responses to antibiotic administration, the formation of H_2_ (Engel et al., 1973; Silverman et al., 2017), and experiments showing that NEC does not develop in germ-free mice (Morowitz et al., 2010; Musemeche et al., 1986). However, while culture-dependent and culture-independent studies have identified differences in fecal microbes between infants with and without NEC, the observed differences have generally varied across studies. Thus, it has been considered unlikely that the presence of a single pathogen or absence of a single commensal is responsible for the disease. A recent meta-analysis of 14 DNA-based studies reported that fecal samples from preterm infants later diagnosed with NEC contained modest but statistically significant increased abundance of facultative anaerobes from the Proteobacteria phylum and a modest decrease in abundance of strict anaerobes (Pammi et al., 2017). As others have previously noted, an understudied approach is to pair taxonomic profiling with functional information about bacterial behavior in response to local conditions within the infant gut (Brown et al., 2018; Pammi et al., 2017).

A longstanding idea has been that inadequate oxygen tension in the intestine, due to reduced regional blood flow (ischemia), may be a key contributor to NEC development (Morowitz et al., 2010; Young et al., 2011). *In vivo* patterns of microbial gene expression in the infant gut may be an indicator of insufficient oxygen supply to the intestine and the progression of NEC. To explore this hypothesis, we retrieved metagenomic and microbial transcriptomic sequence data from prospectively collected time series of samples from four premature infants, two of whom developed NEC (Fig. 1A and Fig. S1A; selected from a larger metagenomic dataset, see (Brown et al., 2018)). The two NEC infants we examined had microbial community compositions with a high abundance of Proteobacteria phyla relative to Firmicutes phyla (Fig. S1C). To probe the response of the gut microbiome to oxygen levels and other aspects of the surrounding conditions, we measured transcript abundances for specific informative set of genes, and compared them to transcript abundances of genes expected to have the opposite response to these conditions (Fig. S1B). Transcript abundance ratios were calculated for each genome at each time point, eliminating biases related to genome relative abundances (See methods in supplemental). This method was chosen given the hypothesis-driven approach taken here, and is distinct from more usual methods for meta-transcriptomic observations.

**Figure 1:**
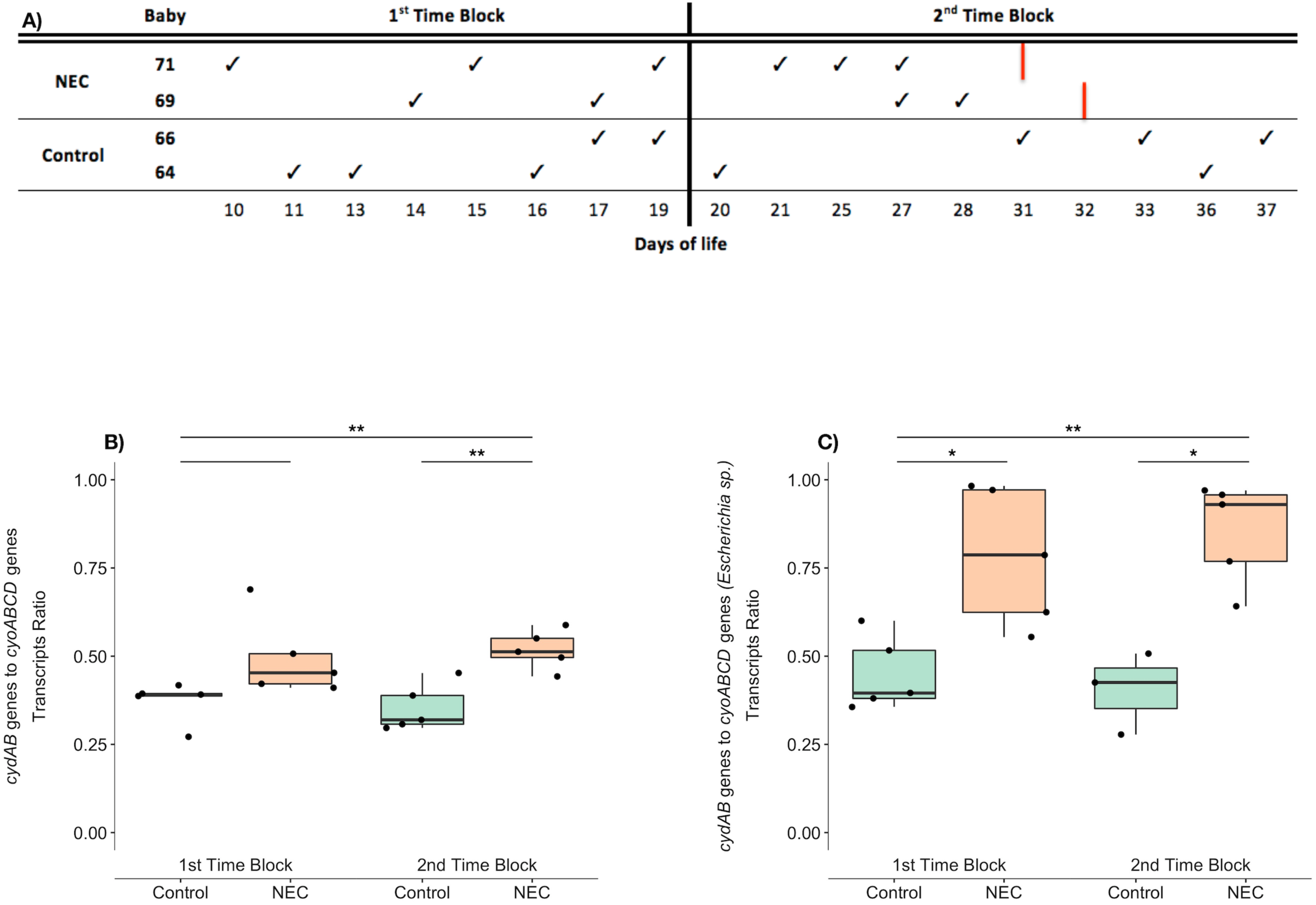
Sampling scheme and microbial transcriptional response to oxygen by gut microbiome of NEC and control premature infants. A. Feces sampling scheme: check mark indicate which infant and at what day of life feces were taken. Red lines on the NEC infant’s rows indicate the day of life (DOL) NEC was detected. DOL10-19 were collapsed into the 1^st^ time block and DOL 20-37 into the 2^nd^ time block.
B. Box plots showing the ratios of *cydAB* and *cyoABCD* transcript abundances averaged over all genomes possessing both sets of genes. Transcript ratios were compared between NEC and control infants in each time block (short lines above) and across all time points (longer lines). Distributions of transcript ratios were compared using Welch’s t-test with Bonferroni correction. Asterisks and double asterisks (^∗^, ^∗∗^) represent*p* < 0.05 and *p* < 0.01, respectively.
C. Box plots showing the ratios of *cydAB* and *cyoABCD* transcript abundances for genomes of bacteria of the genus *Escherichia.* For statistical methods, see (B).

Cytochrome oxidases are complexes of the electron transport chain that pass electrons to O_2_, during aerobic respiration. We examined the expression of bacterial cytochrome oxidase genes with different affinities for oxygen: cytochrome bd oxidase (*cydAB*) with high affinity for oxygen and cytochrome o oxidase (*cyoABCD*) with low affinity for oxygen (D’Mello et al., 1995, 1996). Consistent with these biochemical predictions, a previous study showed that under microaerophilic conditions, there is higher expression of *cydAB* whereas under aerobic conditions expression of *cyoABCD* genes increases (Tseng et al., 1996). In the current study we found statistically higher ratios of *cydAB* relative to *cyoABCD* transcripts in gut microbiomes of infants later diagnosed with NEC compared to the infants that did not develop NEC (Fig. 1B). This observation holds when transcript ratios are averaged for all microorganisms with these genes in each sample, and is significant when comparing across all time points (p = 0.00134). Organisms of the genus *Escherichia* were examined separately, as these organisms were present in all samples and most time points. These organisms also showed higher *cydAB* to *cyoABCD* transcript ratio in the gut of NEC infants compared to control infants (Fig. 1C), and these differences were significant across all time points (p = 0.00006). There are two interpretations for this result. First, bacteria in the guts of the infants that developed NEC may have had lower oxygen availability compared to those that did not develop NEC. Alternatively, the *cydAB* may be more highly expressed than the *cyoABCD* oxidase due to its function in protection against oxidative stress (Giuffrè et al., 2014). For example, the *cyoABCD* expression may have not changed substantially but *cydAB* expression was up-regulated in response to production of radical oxygen species in mitochondria (Poyton et al., 2009), which is produced in response to hypoxia.

To discern between these two explanations for the cytochrome oxidases expression patterns, we examined overall transcription levels of catalase-peroxidase (*katG*), a bacterial gene that encodes an enzyme known to detoxify radical oxygen species (Storz et al., 1990). Importantly, we found no significant differences in *katG* expression between NEC and control samples (Fig. S2A). Furthermore, on an organism-by-organism basis, we examined the transcription levels for *cydAB* and *cyoABCD* relative to known house-keeping genes (Rocha et al., 2015) in both NEC and control infants. Expression of *cydAB* was comparable in both cases, whereas *cyoABCD* significantly varied between NEC and control infants (Fig. S2B), indicating that changes in the transcript ratio of these genes is more likely linked to low O_2_ availability rather than an increase in oxidative stress.

Another potential indicator of conditions in the infant gut is the availability of nitric oxide (NO). Historically, NO has been related to NEC because NO is produced as an antimicrobial during inflammatory response by gut epithelial cells and as it plays a role in controlling blood flow in mesenteric vasculature (Ford et al., 1997; Nowicki et al., 2007). The microbial genetic system we used for examining microbial transcriptional response to NO is the *norVW* genes, coding for NO detoxifying enzymes, and their oppositely transcribed regulating gene *norR* (Jarboe et al., 2010). Nitric oxide binds to constitutively expressed NorR and up-regulates the transcription of *norVW* and down-regulates *norR* expression (Jarboe et al., 2010). Thus, to assess microbial transcriptional response to NO prior to NEC diagnosis, we measured the ratio of transcript abundances for *norVW* and *norR.* Higher relative transcriptional levels of NO detoxifying enzymes compared to *norR* were found in the guts of the control infants across all microbial community members with NO detoxification machinery (Fig. 2A), as well as for the near ubiquitous *Escherichia sp.* (Fig. 2B).

**Figure 2:**
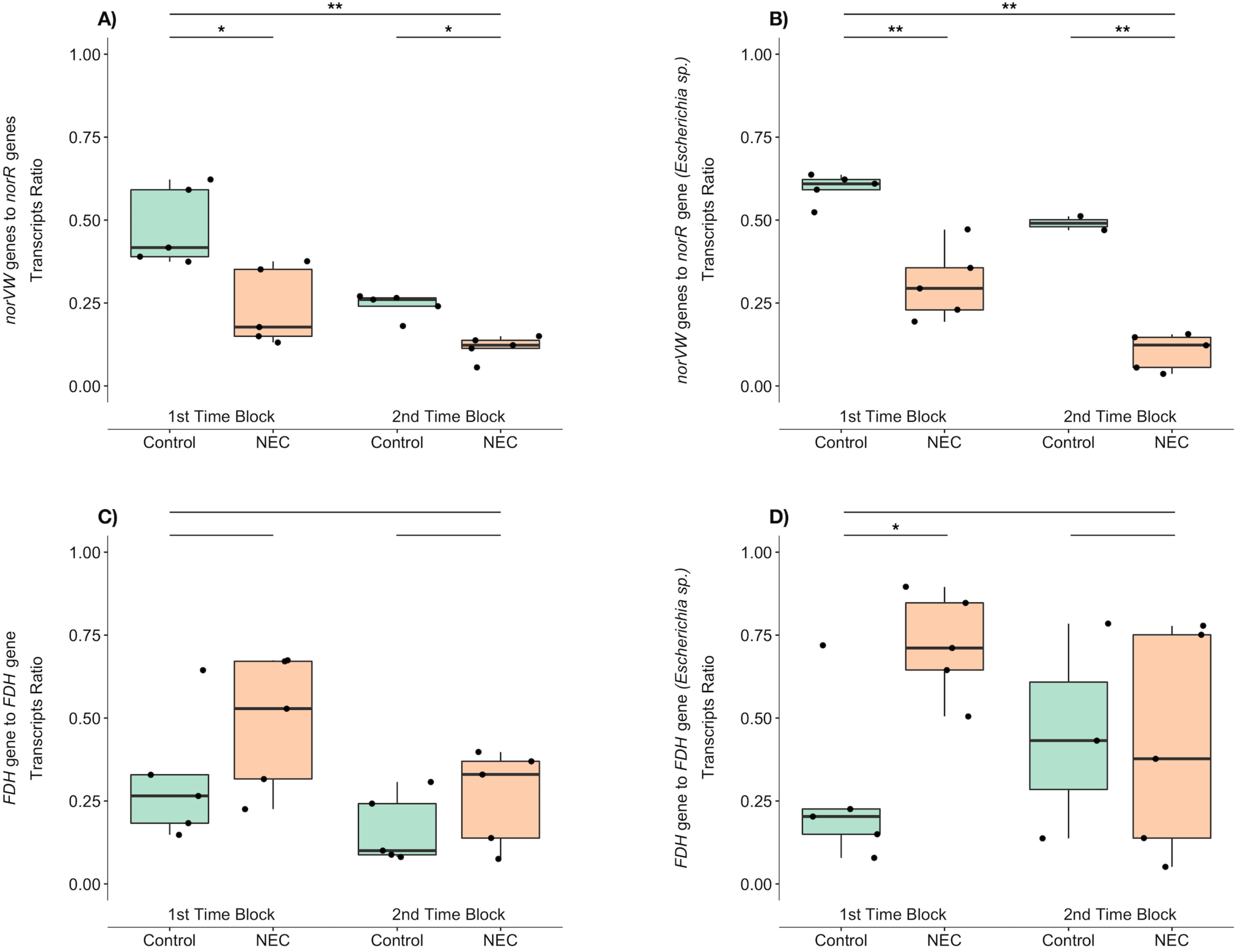
Microbial transcriptional responses to nitric oxide and H_2_ production by gut microbiomes of NEC and control premature infants. A. Box plots showing the ratios of *norVW* and *norR* transcript abundances averaged over all genomes possessing both set of genes. Distributions of transcript ratios were compared using non-parametric Wilcoxon rank-sum test with Bonferroni correction. Asterisks and double asterisks (^∗^, ^∗∗^) represent*p* < 0.05 and*p* < 0.01, respectively.
B. Box plots showing the ratios of *norVW* and *norR* transcript abundances for genomes of the genus *Escherichia.* For statistical methods, see Figure 1B caption.
C. Box plots showing transcript ratios between *FDH* and *FDO* averaged between all genomes possessing both set of genes. For statistical methods, see Figure 1B caption.
D. Box plots showing the ratios of *FDH* and *FDO* transcript abundances in genomes of the genus *Escherichia.* Distributions of transcript ratio were compared using Welch’s t-test (on log10 transformed data) with Bonferroni correction. Asterisks and double asterisks (^∗^, ^∗∗^) represent*p* < 0.05 and *p* < 0.01, respectively.

Interestingly, the ratio of *norVW*to *norR* transcript abundances was higher at earlier compared to later time points, for both NEC and control infants (Welch’s t test, p=0.0089). This probably reflects the response of the preterm infant gut to burgeoning microbial colonizers.

The ability to produce H_2_ is another microbial metabolism that could be an indicator of oxygen levels in the premature infant gut. H_2_ gas, presumably of microbial origin, was previously identified in mural cysts of NEC afflicted infants (Engel et al., 1973). This observation led to development of a method for early detection of NEC based on breath hydrogen excretion test. However, this test did not become widely accepted due to its low positive prediction value (Cheu et al., 1989). A key source of microbial H_2_ production is formate dehydrogenase (FDH), which converts excess formate formed during mixed acid fermentation into CO_2_ and H_2_ (Sawers, 2005). There are three different kinds of formate dehydrogenase’s that are expressed under different conditions, but only one type produces H_2_ (Sawers, 2005). Two of the three formate dehydrogenases were found in most genomes. Thus, we compared transcription of FDH genes, which are responsible for formate removal from the cell and H_2_ production, to the transcript levels of FDO genes that convert formate to H^+^ and CO_2_ without production of H_2_. The average FDH/FDO transcript ratio of the whole microbial community was not significantly different between NEC and control infants (Fig. 2C). We also compared transcript ratios for members of the genus *Escherichia* and found higher FDH compared to FDO expression in NEC diagnosed infants compered to control infants (Fig. 2D). Interestingly, significantly higher expression occurred at earlier time points in NEC compared to control cases.

Given the findings related to H_2_ production, we investigated genome–resolved metagenomic data from five infants that developed NEC and seven that did not develop NEC for whom transcriptomic data were not available. We identified genomes encoding FDH and auxiliary genes and found that bacteria with the genetic capability to produce H_2_ occurred at higher relative abundances in gut microbiomes of infants that developed NEC. These bacteria were significantly more abundant in NEC infant compared to control infant gut microbiomes in samples taken at 20-40 DOL (Day of life), in both the smaller (Fig. S2C) and the larger meta-genomic data sets (Fig. S2B). Although significant, the results are very variable. Interestingly, transcript abundance data indicate capacity for higher H_2_ production at earlier time points (DOL 10-20) in the infants that developed NEC (Fig. 2D). This finding underlines the importance of distinguishing genetic capacity from activity.

In combination, information about O_2_ availability and H_2_ production by microbial communities suggest the onset of hypoxic conditions prior to NEC formation in the gut of premature infants. Pig models have shown that fetuses and premature neonates have higher blood flow to the gut than in more developed gut systems (Nankervis et al., 2001), and this may lead to more oxygenated conditions in the gut lumen. This is consistent with previous observations showing that the gut of premature infants is initially colonized with more aerobic organisms that later shifts to become more anaerobic communities (Brooks et al., 2015; La Rosa et al., 2014; Sharon et al., 2013). During NEC, however, hypoxic conditions have been observed in the gut tissue of many patients (Chen et al., 2016) and histologic examination of removed dead tissue demonstrates coagulation necrosis, evidence for ischemic injury (Ballance et al., 1990). Yet, hypoxia is highly disputed as a primary controlling factor of NEC (Chen et al., 2016; Crissinger, 1994; Neu, 2005; Nowicki and Nankervis, 1994; Young et al., 2011). Recent theories on NEC development point to the role of gut tissue immaturity, exhibiting high expression of Toll-like receptor 4 (TLR4), an innate immune receptor that recognizes lipopolysaccharides on the surfaces of Gram-negative bacteria. Activation of TLR4 initiates an inflammatory response, which can impair intestinal microcirculation and oxygen delivery (Yazji et al., 2013). These phenomena may explain the observed transcriptional response to lower O_2_ levels in the gut microbiomes of NEC infants (Fig. 1BC). This circumstance is distinct from that that occur in the mature gut, where low O_2_ is linked to a healthy condition and aerobicity is linked to inflammation (Litvak et al., 2017).

Increased expression of inducible nitric oxide synthase (iNOS) in gut epithelial cells often occurs as a part of the inflammatory response, and recent studies have suggested that TLR4-mediated iNOS expression is a key element of NEC progression (Jilling et al., 2006). Due to the short half-life of NO and the close proximity of epithelial cells to the gut lumen, this is probably the main source of NO in the gut lumen (Singer et al., 1996). Higher expression of iNOS in host epithelial cells was found during NEC, both in mice and in gut tissue removed from NEC patients by surgery (Ford et al., 1997; Jilling et al., 2006). Counter-intuitively, in this study we found down regulation of bacterial genes for NO detoxification in the gut of NEC infants (Fig. 2AB), indicating lower NO levels in the gut lumen.

Resolving these seemingly conflicting observations may require a more careful examination of the factors that regulate host iNOS activity. The reaction that is catalyzed by nitric oxide synthases is the conversion of L-arginine and oxygen into L-citrulline and NO (Bogdan, 2015). Thus, Oxygen levels are a controlling factor for iNOS activity (Bogdan, 2015). We suggest that in situations where O_2_ supply is reduced, the iNOS response in gut epithelial cells might be frustrated and the antimicrobial response is largely inactivated. Previously, a review paper also proposed that although hypoxic conditions and inflammation will induce expression of iNOS gene, hypoxia would reduce NO production by the iNOS enzyme (Robinson et al., 2011). However, we cannot rule out the possibility that lower availability of another iNOS substrate, L-arginine, may limit the activity of this enzyme during NEC development, as low L-arginine concentrations are associated with NEC (Watkins and Besner, 2013).

In conclusion, this study shows that meta-transcriptomic data can probe physiological conditions that gut microbial communities experience. Despite the small sample size, we demonstrated that the transcript abundance of H_2_ forming formate dehydrogenase, relative to non-H_2_ forming formate dehydrogenase, is significantly higher at early time points for *Escherichia sp.* genomes in infants that develop NEC. The higher transcriptional response to NO by gut microbiome in control compared to NEC infants was even more significant. We suggest that lower NO concentrations in infants that develop NEC are probably related to lower O_2_ availability, due to reduced blood flow that restricts NO synthesis in epithelial gut cells. Lower O_2_ availability also increased microbial transcript ratios for the cytochrome oxidase that evolved to function in low O_2_ environments relative to the cytochrome oxidase optimized for high O_2_ conditions in NEC infants compared to control infants. A larger study is needed to verify these results and confirm our main conclusion, that the gut microbiome, in early stages of NEC development, senses and responds to different physiological conditions compared to microbiomes in guts where NEC is not developing. Understanding the physiological conditions during early stages of NEC development may inspire the development of new tools for early NEC diagnosis, enabling treatment prior to the fast deterioration phase of the disease.

## Acknowledgments

This research was supported by the National Institutes of Health (NIH) under award no. RAI092531A. This work was supported in part by the March of Dimes Foundation research Grant no. 5-FY10-103 (to MJM). YS work was supported by postdoctoral fellowship grant no. 2016-67012-24717 from the USDA National Institute of Food and Agriculture.

